# Dysconnection and cognition in schizophrenia: a spectral dynamic causal modeling study

**DOI:** 10.1101/2022.10.09.511459

**Authors:** Tahereh S. Zarghami, Peter Zeidman, Adeel Razi, Fariba Bahrami, Gholam-Ali Hossein-Zadeh

**Affiliations:** Bio-Electric Department, School of Electrical and Computer Engineering, College of Engineering, University of Teran, Tehran, Iran; Human Motor Control and Computational Neuroscience Laboratory, School of Electrical and Computer Engineering, College of Engineering, University of Tehran, Tehran, Iran; The Wellcome Centre for Human Neuroimaging, University College London, Queen Square, London WC1N 3AR, UK; Turner Institute for Brain and Mental Health, Monash University, Clayton, VIC; Monash Biomedical Imaging, Monash University, Clayton, VIC; CIFAR Azrieli Global Scholars Program, CIFAR, Toronto, Canada

**Keywords:** Effective Connectivity, dynamic causal modeling, resting state fMRI, schizophrenia, dysconnection hypothesis, cognitive impairment, canonical correlation analysis

## Abstract

Schizophrenia (SZ) is a severe mental disorder characterized by failure of functional integration (aka dysconnection) across the brain. Recent functional connectivity (FC) studies have adopted functional parcellations to define subnetworks of large-scale networks, and to characterize the (dys)connection between them, in normal and clinical populations. While FC examines statistical dependencies between observations, model-based effective connectivity (EC) can disclose the causal influences that underwrite the observed dependencies. In this study, we investigated resting state EC between the subnetworks of seven large-scale networks, in 66 SZ and 74 healthy subjects from a public dataset. The results showed that a remarkable 33% of the effective connections (among subnetworks) of the cognitive control network had been pathologically modulated in SZ. Further dysconnection was identified within the visual, default mode and sensorimotor networks of SZ subjects, with 24%, 20% and 11% aberrant couplings. Overall, the proportion of diagnostic connections was remarkably larger in EC (24%) than FC (1%) analysis. Subsequently, to study the neural correlates of impaired cognition in SZ, we conducted a canonical correlation analysis between the EC parameters and the cognitive scores of the patients. As such, the self-inhibitions of supplementary motor area and paracentral lobule (in the sensorimotor network) and the excitatory connection from parahippocampal gyrus to inferior temporal gyrus (in the cognitive control network) were significantly correlated with the social cognition, reasoning/problem solving and working memory capabilities of the patients. Future research can investigate the potential of whole-brain EC as a biomarker for diagnosis of brain disorders and for cognitive assessment.

## 1 Introduction

Schizophrenia (SZ) is a debilitating brain disorder characterized by episodes of psychosis; common symptoms include delusions, hallucinations, disorganized thinking, social withdrawal and apathy. SZ is also associated with a wide range of cognitive impairments, spanning from basic perceptual processes to complex nonsocial and social cognitive functions (Green et al., 2019). The dysconnection hypothesis (Friston and Frith, 1995; Friston et al., 2016a) tries to bridge the explanatory gap between the symptoms and signs of SZ and the underlying neuronal pathophysiology. To this end, it casts psychosis as “aberrant neuromodulation of synaptic efficacy that mediates the (context-sensitive) influence of intrinsic and extrinsic (long-range) connectivity” (Friston et al., 2016a).

Among different neuroimaging modalities, functional magnetic resonance imaging (fMRI) predominates in system-level connectomic studies. Particularly, the task-free version - known as resting state fMRI - is better tolerated by clinical populations, and circumvents the need for stringent subject compliance. Today, resting state fMRI features are being rigorously examined to identify potential biomarkers for diagnosis and prognosis of different brain disorders (Abraham et al., 2017; Chen et al., 2022; Damoiseaux, 2012; Drysdale et al., 2017; Franzmeier et al., 2017; Hohenfeld et al., 2018; Khalili-Mahani et al., 2017; Pfannmöller and Lotze, 2019; Rashid and Calhoun, 2020; Taylor et al., 2021; Viviano et al., 2018; Wang et al., 2018).

Earlier connectomic studies have investigated the (dys)connection between different *nodes* of a network, defined based on local *anatomical* demarcations (Bassett et al., 2008; Liang et al., 2006; Supekar et al., 2008; Zhou et al., 2007). More recently, *functional* parcellation of large-scale networks has gained interest (Allen et al., 2014; Glasser et al., 2016; Gordon et al., 2014; Schaefer et al., 2018; Tian et al., 2020), furnishing insight into the interaction of finer-scale functional patterns as subcomponents of larger-scale distributed networks. Empirical evidence from numerous functional connectivity (FC) studies suggests that the interaction of constituent subnetworks of intrinsic networks has been disrupted in SZ. Dysconnection of subnetworks (within and across principal modes^1^/networks) has been associated with the clinical and genomic characteristics of SZ patients in several FC studies (Du et al., 2020; Miller et al., 2016; Rabany et al., 2019; Rashid et al., 2019).

However, since FC is meant to quantify *statistical dependencies* between the observations (i.e. neurophysiological recordings), it does not reveal the directed *effective/causal influences* that underwrite these dependencies (Friston, 2011). The latter is referred to as effective connectivity (EC). In EC analysis, a generative model should be specified, which can predict the observations based on a biophysically-grounded model of the network dynamics, called a dynamic causal model (DCM) (Friston et al., 2003). Given some empirical data, the parameters of this model would be estimated such that the model optimally explains the observations—in a procedure known as model inversion (Friston et al., 2007; Zeidman et al., 2022). The most established EC model, in the context of resting state fMRI, is called spectral DCM (Friston et al., 2014a; Razi et al., 2015). The term *spectral* highlights the nature of the observations (i.e., cross spectral densities between signals) that the model is designed to explain. Notably, cross spectrum is the Fourier counterpart of cross-covariance function, which (at zero time-lag and normalized) is the most conventional FC measure. In other words, spectral DCM is a generative model of how FC is realized. Notably, the effort that goes into the specification of a generative model and performing model inversion in EC, results in the separation of neuronal-level coupling from observation-level dependencies.

We speculated that resting state EC among the subnetworks of the principal intrinsic networks of the brain is disrupted in SZ. To test this hypothesis, we identified 50 subnetworks of seven large-scale networks of the brain, using constrained spatial independent components analysis (CSICA), in 74 HC and 66 SZ subjects. The networks comprised the: subcortical (SC), auditory (AUD), sensorimotor (SM), visual (VIS), cognitive control (COG), default mode network (DMN) and cerebellum (CB). Subject-level estimates of EC (using spectral DCM) were analyzed, for each network separately, in a parametric empirical Bayesian (PEB) scheme, to identify group differences that characterize the conjectured dysconnection within the examined networks. Additionally, the same networks were analyzed using FC, for comparison. We asked whether the two (FC and EC) approaches reveal different aspects of network dysconnection in SZ. We also asked which large-scale networks - and to what extent – are dysconnected in this disorder.

In the second part of the research, we investigated the neural correlates of cognitive impairment in SZ. Schizophrenia has been associated with a wide range of cognitive deficits including aberrations in speed of processing, attention, working memory, verbal and visual learning, problem solving and social cognition (Green et al., 2019). Cognitive impairment is a core feature of SZ, and the prime driver of severe disabilities in functional outcomes (including occupational, social, and economic performance) of the patients (Green, 2006; Green et al., 2000; Lepage et al., 2014; Tripathi et al., 2018). These impairments cannot be explained by the positive symptoms of the disorder or the medication effects (Green et al., 2019), and they are largely unresponsive to antipsychotic treatment (Tripathi et al., 2018). Nevertheless, the neural basis of cognitive impairment in SZ is still poorly understood (Alkan et al., 2021).

In order to establish functional validity for the EC estimates, and to elucidate the neural associates of cognitive impairment in SZ, we conducted a canonical correlation analysis (CCA) between the EC parameters and the cognitive scores of the patients (from MCCB^2^ tests). To achieve a stable CCA model, we observed the practical recommendations (about sufficient sample to feature ratio) from recent technical reports (Helmer et al., 2020; Yang et al., 2021) using appropriate feature selection procedures. From this analysis, we asked which effective connections and cognitive traits are mostly correlated in the patients, and whether such a linear association holds some degree of generalizability beyond the current sample.

## 2 Materials and Methods

### 2.1 Dataset and pre-processing

We analyzed the publicly available schizophrenia dataset of the Center for Biomedical Research Excellence (COBRE)^3^ (Çetin et al., 2014), which includes 72 SZ patients and 76 healthy control (HC) subjects (18-65 years old). The patients had been diagnosed using the Structured Clinical Interview for DSM-IV^4^ Axis I Disorders (SCID-I) (First et al., 2002), and were receiving antipsychotic medications. Five-minute resting state scans were acquired on a 3-Tesla Siemens Tim Trio scanner, during which subjects were instructed to keep their eyes open and fixate on a central cross. A total of 150 (T2*-weighted) functional volumes were collected using a gradient-echo EPI sequence, with the following settings: TR = 2 s, TE = 29 ms, flip angle = 75°, 33 axial slices, ascending acquisition, matrix size = 64 × 64, voxel size = 3.75 × 3.75 × 4.55 mm, field of view = 240 mm. A high-resolution T1-weighted structural image had also been collected for each subject.

Standard pre-processing of functional data was performed using the SPM12 software^5^. In brief, the first 5 volumes were discarded to allow for T1 equilibration; the remaining images were realigned to the first volume (for motion correction), slice-timing corrected, co-registered to the anatomical image of the corresponding subject, warped to the standard Montreal Neurological Institute (MNI) template (Collins et al., 1998), resampled to 3 *mm*^3^ isotropic voxels, and smoothed with a Gaussian kernel (FWHM = 6 mm).

Two disenrolled subjects, one subject with incomplete functional session and five subjects with excessive head motion (translation > 4 mm or rotation > 4 degrees) were excluded. For the remaining subjects, the mean framewise displacement (FD) was reasonably low (0.41 ± 0.22 mm) (Power et al., 2012). The demographics of the retained 140 subjects (74 HC/66 SZ) have been summarized in Table 1. There were no significant differences in age, gender proportion, maximum head translation and rotation between the two groups (uncorrected p-values = 0.12, 0.26, 0.13, 0.36, respectively). The patients were receiving antipsychotic medications, which were converted to their chlorpromazine (CPZ) equivalents.

**Table 1:** Demographics of the participants.

|  | Number | Female /Male | Age | Max head translation (mm) | Max head rotation (degrees) | CPZ equivalent (mg/day) |
| --- | --- | --- | --- | --- | --- | --- |
| **Healthy Controls** | 74 | 23/51 | 35.8 ± 11.6 | 0.94 ± 0.65 | 0.83 ± 0.53 | - |
| **Schizophrenia Patients** | 66 | 13/53 | 38.3 ± 14.2 | 1.12 ± 0.76 | 0.93 ± 0.73 | 363.1 ± 305.0 |

CPZ: Chlorpromazine

### 2.2 Cognitive profile

The MATRICS^6^ consensus cognitive battery (MCCB) tests (August et al., 2012; Kern et al., 2011; Nuechterlein et al., 2008) been conducted to estimate the cognitive performance of the participants in seven domains: speed of processing, attention/vigilance, working memory, verbal learning, visual learning, reasoning/problem solving, and social cognition. The composite MCCB score had been calculated as the standardized sum of all seven domains, based upon published normative data (Kern et al., 2011). Out of the 140 subjects included in our analysis, MCCB T-scores were available for a total of 117 participants (59 HC, 58 SZ). We compared the performance of HC and SZ subjects in each domain and in their composite scores, using two-sample permutation-based t-tests with maxT correction for multiple comparisons (Westfall and Young, 1993).

### 2.3 Network identification

We conducted a refined functional parcellation of the brain, per subject, using (spatially) constrained spatial ICA (CSICA) (Lin et al., 2010), as implemented in the Group ICA of fMRI Toolbox (GIFT^7^). The spatial constraints were imposed using aggregate functional networks from a previous large group study by (Allen et al., 2014). Notably, CSICA maximizes the independence of spatial components (i.e. networks) for each individual, while acknowledging spatial variability at the subject level and preserving network correspondence at the group level (Lin et al., 2010). As spatial priors, we used the 50 aggregate functional networks identified in (Allen et al., 2014) based on a refined group spatial ICA analysis on 405 subjects^8^. These 50 functional parcels constituted subcomponents of seven reproducible large-scale resting state networks; namely, the subcortical (SC), auditory (AUD), sensorimotor (SM), visual (VIS), cognitive control (COG), default mode network (DMN) and the cerebellum (CB) (Fig. 1-B). The list of subnetworks and their abbreviations is available in Table 2.

**Fig. 1:**
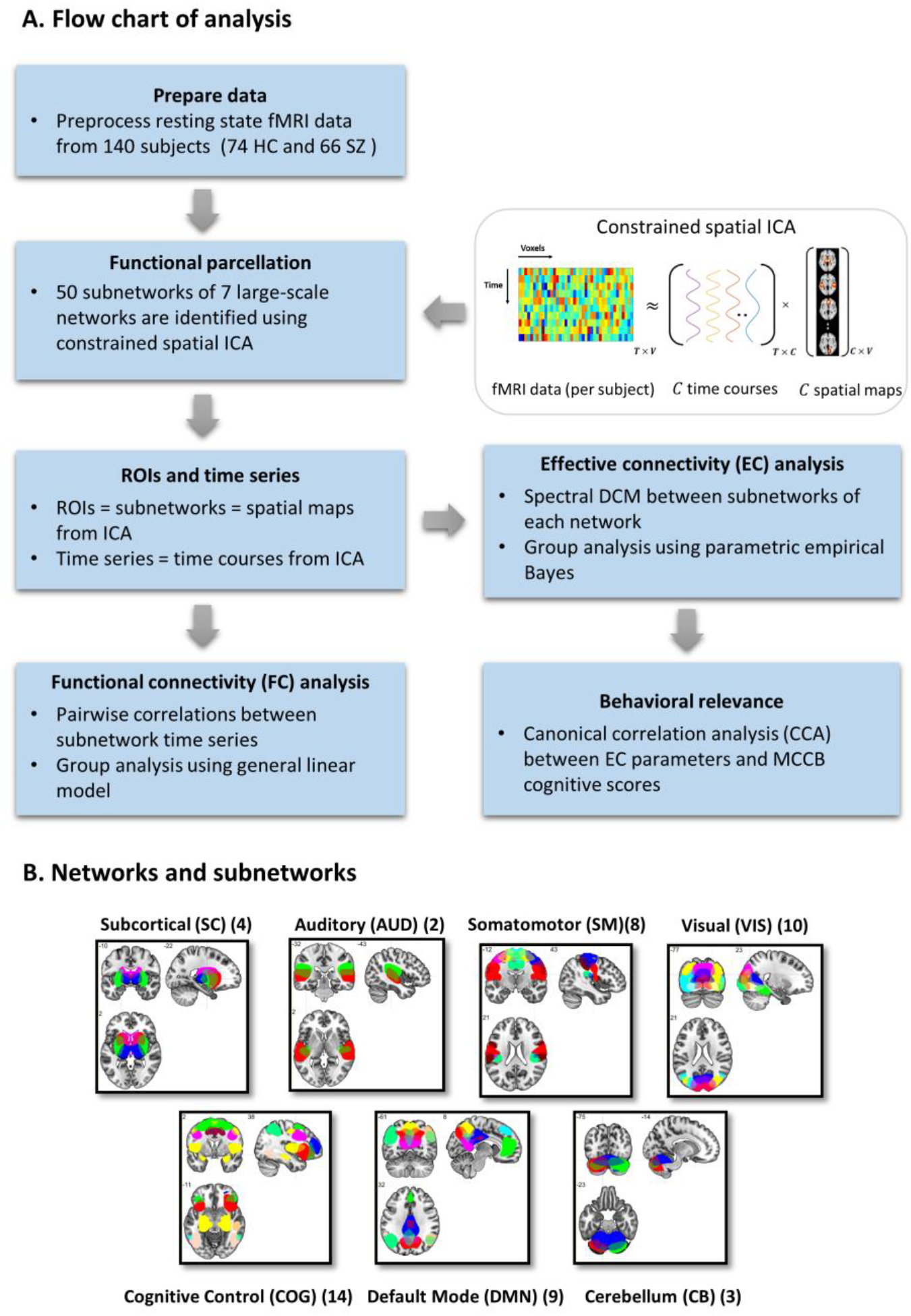
(A) Flowchart of analysis. (B) The 50 subnetworks used as spatial priors belong to seven large-scale resting state networks. The number of constituent subnetworks of each network has been mentioned in parentheses. Subject-specific versions of these subnetworks and their corresponding timeseries were identified using constrained spatial ICA, and adopted as regions of interest (ROIs) and ROI timeseries in functional and effective connectivity analyses. Cognitive relevance of the EC parameters was assessed using canonical correlation analysis. The subnetwork abbreviations are listed in Table 2.

**Table 2:** Names and abbreviations of 50 aggregate subnetworks from (Allen et al., 2014) used herein as spatial priors/templates to derive subject-specific subnetworks using constrained spatial ICA. Each template is a mask containing one or more (typically bilateral) clusters within the brain. The peak coordinates of these templates are available in Table S1 and Fig S2 of (Allen et al., 2014). The group maps have been publicly shared by the authors at https://trendscenter.org/data

| Network | Subnetwork abbreviation | Subnetwork full name |
| --- | --- | --- |
| **Subcortical (SC)** | Putamen1 | Putamen 1 |
|  | Putamen2 | Putamen 2 |
|  | Thalamus | Thalamus |
|  | Caudate | Caudate |
| **Auditory (AUD)** | STG1 | Superior temporal gyrus 1 |
|  | STG2 | Superior temporal gyrus 2 |
| **Sensorimotor (SM)**<br>**(aka somatomotor)** | PreCG | Precentral gyrus |
|  | SPL | Superior parietal lobule |
|  | R-PoCG | Right postcentral gyrus |
|  | L-PoCG | Left postcentral gyrus |
|  | ParaCL1 | Paracentral lobule 1 |
|  | ParaCL2 | Paracentral lobule 2 |
|  | PoCG | Postcentral gyrus |
|  | SMA | Supplementary motor area |
| **Visual (VIS)** | Cuneus | Cuneus |
|  | FFG | Fusiform gyrus |
|  | CalcarineG | Calcarine gyrus |
|  | Cuneus | Cuneus |
|  | SOG | Superior occipital gyrus |
|  | MTG | Middle temporal gyrus |
|  | LingualG | Lingual gyrus |
|  | MOG | Middle occipital gyrus |
|  | R-MOG | Right middle occipital gyrus |
|  | L-MOG | Left middle occipital gyrus |
| **Cognitive Control (COG)** | aInsula | Anterior insula |
|  | RSN-SMA | Supplementary motor area |
|  | MiFG1 | Middle frontal gyrus 1 |
|  | MiFG2 | Middle frontal gyrus 2 |
|  | PreCG | Precentral gyrus |
|  | IPL | Inferior parietal lobule |
|  | R.STG+IFG | Right superior temporal gyrus +<br>Inferior frontal gyrus |
|  | R-IPL | Right inferior parietal lobule |
|  | pInsula | Posterior insula |
|  | L-IPL | Left right inferior parietal lobule |
|  | PHG | Parahippocampal gyrus |
|  | IFG | Inferior frontal gyrus |
|  | MCC | Middle cingulate cortex |
|  | ITG | Inferior temporal gyrus |
| **Default Mode Network (DMN)** | Precuneus1 | Precuneus 1 |
|  | ACC | Anterior cingulate cortex |
|  | PCC1 | Posterior cingulate cortex 1 |
|  | PCC2 | Posterior cingulate cortex 2 |
|  | Precuneus2 | Precuneus 2 |
|  | MiFG+SFG | Middle frontal gyrus + Superior frontal gyrus |
|  | R-AG | Right angular gyrus |
|  | L-AG | Left angular gyrus |
|  | L.MTG+IFG | Left middle temporal gyrus + Inferior frontal gyrus |
| **Cerebellum (CB)** | L-CB | Left cerebellum |
|  | R-CB | Right cerebellum |
|  | CB | Cerebellum |

The time courses corresponding to subject-specific ICs (i.e. subnetworks) were detrended and orthogonalized with respect to the subject’s estimated motion parameters and their first derivatives. The series were further despiked using AFNI’s 3dDespike algorithm, which detects outlier time points and replaces them with spline interpolations. The subject-specific subnetworks and their post-processed time courses were subsequently analyzed using functional and effective connectivity methods.

### 2.4 Functional connectivity analysis

Functional connectivity was computed as the sample covariance matrix of the subnetwork time series for each individual. These FC matrices were then averaged over the subjects of each group (Fig. 3-A). To estimate group differences, we set up a multiple linear regression model for each FC entry as follows: FC (per subject) was the dependent variable; diagnosis was the regressor of interest; age, gender and medication dosage were covariates (Allen et al., 2011; Damaraju et al., 2014). As such, p-values for the significance of diagnosis coefficients and their corresponding t-statistics were recorded. The p-values were adjusted for multiple comparisons based on the false discovery rate (FDR) approach (Benjamini and Hochberg, 1995). The group difference matrices in Fig. 2-B and Fig. 2-C reflect – *sign*(*t* – *statistic*) × log_10_ *q_FDR_* values, for entries with *q_FDR_* < 0.05.

**Fig. 2:**
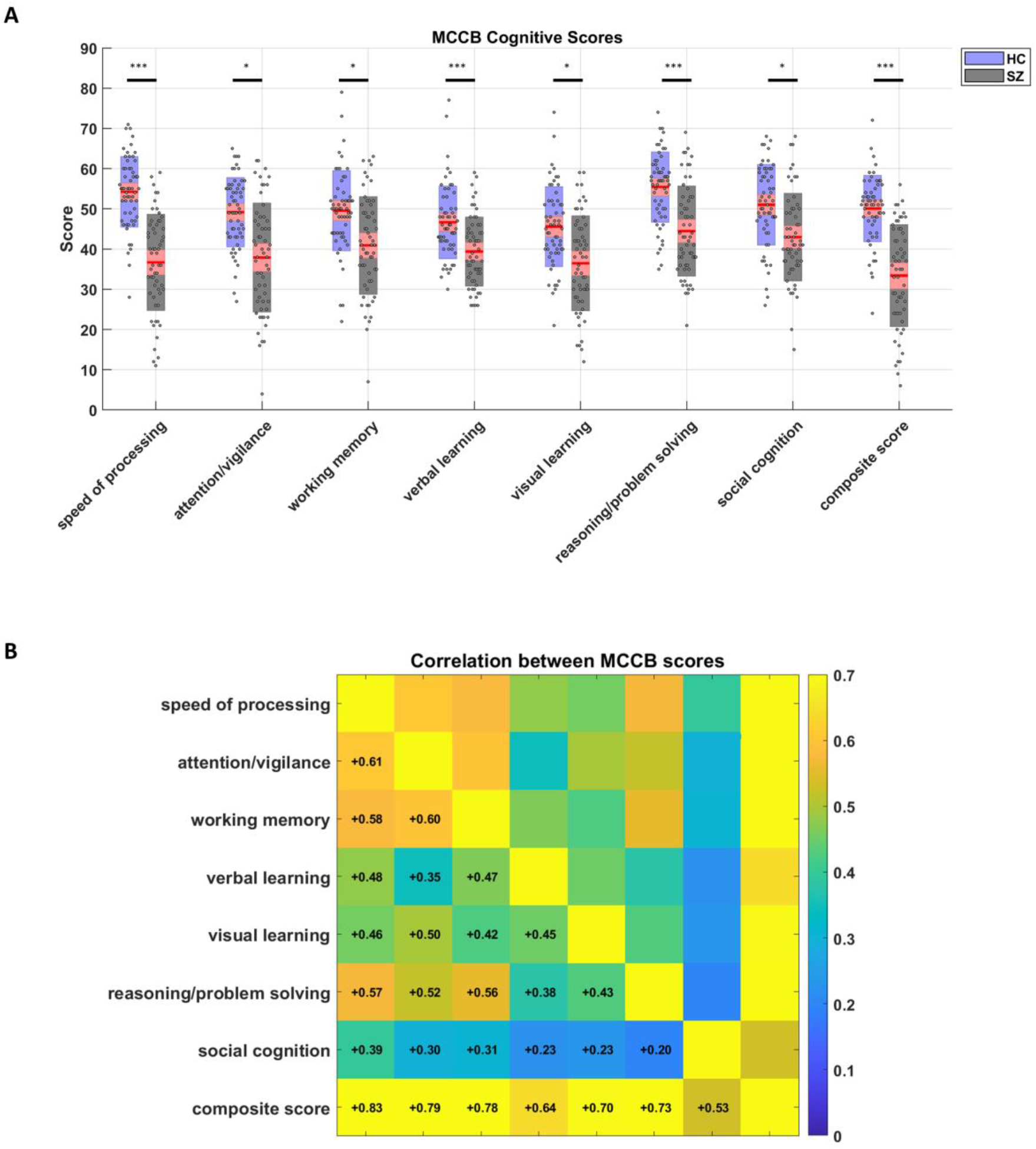
Behavioral (cognitive) results. (A) Statistical comparison of cognitive performance in HC and SZ subjects, as measured by the seven tests of the MCCB and the composite score. The red box plots show 95% confidence intervals around the mean of the scores, while blue/gray boxes mark one STD of the scores for HC/SZ subjects. The standardized MCCB scores are scattered over the boxes. Asterisks indicate statistical significance based on two-sample permutation t-tests with maxT correction for multiple comparisons. *: p <0.05, **: p < 0.01, ***: p < 0.001. (B) Correlation of MCCB scores between different domains, across all subjects.

**Fig. 3:**
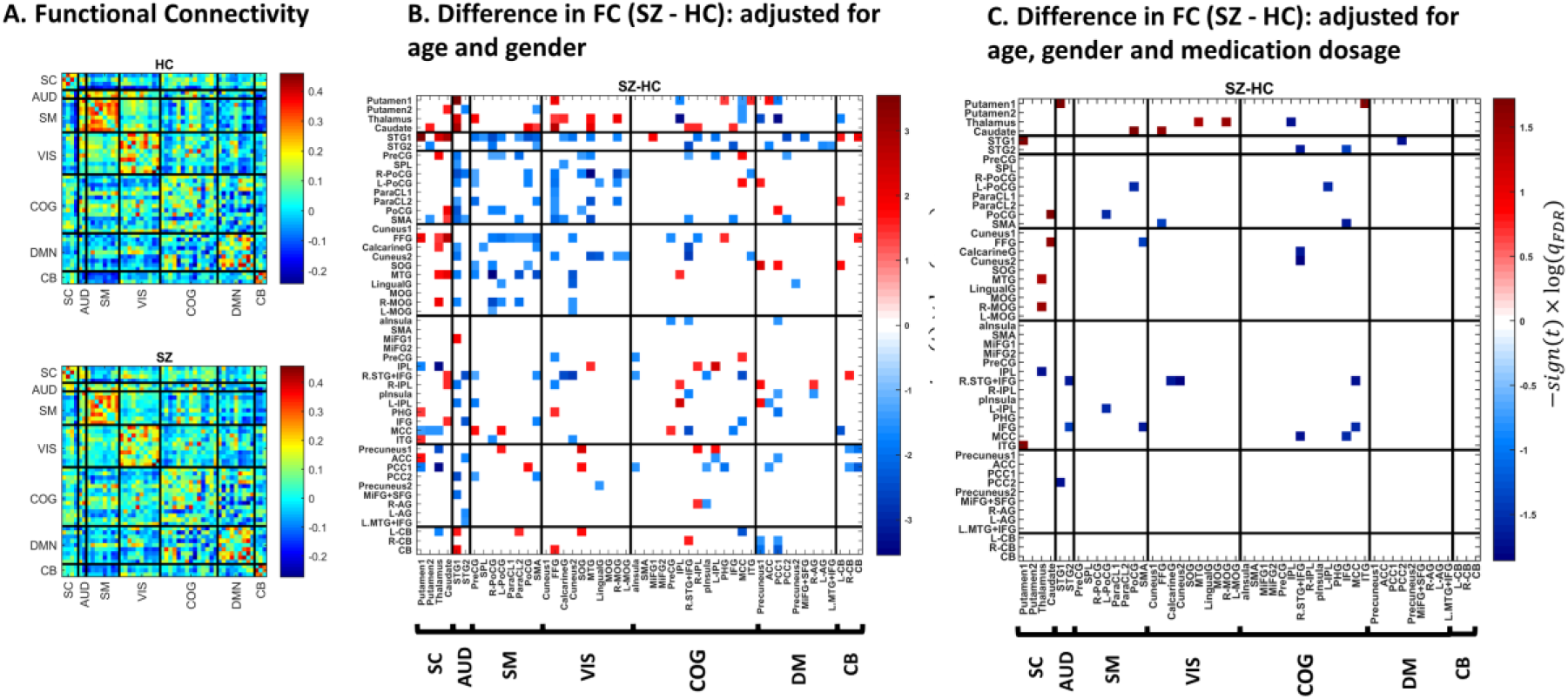
Functional connectivity analysis. (A) Average FC pattern for the HC and SZ group. Diagonal (unity) entries have been removed for better visualization. (B) Difference in FC (SZ - HC), adjusted for age and gender. The colorbar denotes – *sign*(*t*) × log (*q_FDR_*), where the t-statistic is computed from the estimated coefficient in a multiple linear regression model that includes one FC entry (across subjects) as the dependent variable, diagnosis label as the independent variable, plus age and gender as covariates. P-values corresponding to the coefficients’ t-stats have been corrected using the FDR method, and the entries corresponding to *q_FDR_* < 0.05 are marked in color. The 50 sub-labels reflect the constituent subnetworks of seven large-scale networks. (C) Difference in FC (SZ - HC), adjusted for (CPZ equivalent of) medication dosage, in addition to age and gender. Network and subnetwork abbreviations are available in Table 2.

### 2.5 Effective connectivity analysis

Effective connectivity analysis for resting state fMRI was conducted using spectral DCM (Friston et al., 2014a). Group effects were modeled using a hierarchical Bayesian framework known as parametric empirical Bayes (PEB) (Friston et al., 2015; Friston et al., 2016b), as described later.

#### 2.5.1 Spectral dynamic causal model

Spectral DCM was used to estimate the effective connections within each of the seven large-scale resting state networks. Briefly, spectral DCM specifies how complex cross spectra of the BOLD signals are generated from regional hemodynamic responses to the neuronal dynamics of a (biophysically plausible and endogenously driven) neural network. Under local linearity assumptions on the dynamical system, together with parametrized power-law distributions for the noise cross spectra, the model admits a deterministic form that can be efficiently inverted in spectral domain (Friston et al., 2014a; Razi et al., 2015).

The generative model of spectral DCM was then fitted to the cross spectral densities estimated from empirical data using multivariate autoregressive models (as implemented in spm_dcm_fMRI_csd in SPM12). In this routine, the optimal^9^ model parameters are estimated by maximizing a variational lower bound, called free energy^10^, on the log Bayesian model evidence (Friston et al., 2007; Zeidman et al., 2022). Thereafter, maximized free energy is used for model comparison, and the parameter posterior distributions are used for inference about the effective connections.

To estimate the EC among subnetworks of each of the seven large-scale networks, a fully connected network structure was initially assumed for each subject, per network. This assumption was later refined iteratively by incorporating group information (as empirical priors) on the subject-level connections. These empirical priors were computed in a Bayesian framework that facilitates group inference for dynamic causal models (Friston et al., 2015; Friston et al., 2016b; Zeidman et al., 2019)—as explained next.

#### 2.5.2 Group analysis using parametric empirical Bayes

PEB is a hierarchical Bayesian model, particularly useful for estimating group effects in DCM studies. Operationally, PEB is a Bayesian general linear model (GLM) that partitions between-subject variability into designed group effects (such as group mean and difference) and additive random effects. It can also be regarded as a generalization to the *summary statistic* random effects approach, with the advantage that PEB takes the full posterior densities of the first level (i.e. DCM) parameters to the second (between-subject) level. That is, PEB accounts for the posterior uncertainty of the parameters in addition to their expected (point estimate) values. Mathematically, the PEB model is specified as:

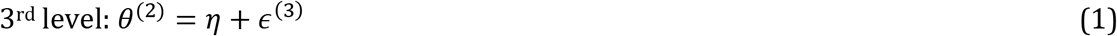

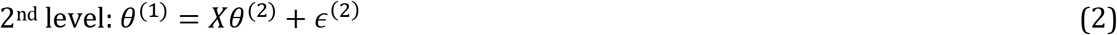

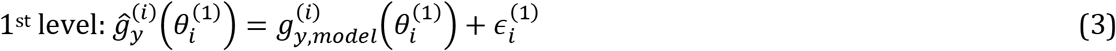

In Eq. 3, the observed (cross-spectral) data features from subject *i* are modeled as having been generated by a spectral DCM with parameters 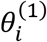 and sampling error 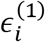. DCM parameters *θ*^(1)^ are themselves described by a GLM in Eq. 2, with design matrix *X,* group parameters *θ*^(2)^ and between-subject variability *ϵ*^(2)^. The columns of X encode the hypothesized sources of inter-subject variability (e.g., group mean and difference) while *ϵ*^(2)^ accounts for the random effects. Hence, each corresponding entry in *θ*^(2)^ is the group-level effect of one covariate on one connection. These group effects, in turn, have priors specified at the 3^rd^ level, as noted in Eq. 1. The parameters and noise components are assumed to be normally distributed, and estimated through an iterative variational Laplace scheme (Friston et al., 2007; Zeidman et al., 2022).

Specifically, the second level (group) parameters are estimated by assimilating the posteriors of the first level parameters. These group parameters then serve as empirical priors for estimation at the first level. And this iterative scheme continues until convergence. Notably, the empirical priors are ‘typical’ group values that guide subject-level inferences and circumvent local maxima problems (Friston et al., 2015; Zeidman et al., 2019).

In summary, first we used spectral DCM to infer effective connectivity for each subject. Then the hierarchical model of PEB was used to integrate subject-level results for group analysis. The design matrix of PEB (X in Eq. 2) comprised of five columns/regressors: a constant column (of 1’s) to capture the commonalities across all participants; a second column to encode group differences between SZ and HC subjects (encoded with +1’s and −1’s); and three other columns containing age, gender and (CPZ equivalent of) antipsychotic medication dosage, as covariates. Similarly, two other PEB models were set up to capture group-specific average patterns of EC.

In each GLM, all DCM parameters were initially allowed to contribute to all group effects. This assumption was refined post hoc using exploratory Bayesian model reduction (BMR) on the group-level posteriors. That is, parameters that did not contribute to the model evidence were recursively pruned from the parent (full) model to generate reduced models. Usually, no single reduced model is the overall winner with probability greater than 0.95. Hence, Bayesian model averaging (BMA) was used to compute the weighted average of each parameter across the top 256 reduced models, where the weights corresponded to the posterior probabilities of these models (Hoeting et al., 1999; Penny et al., 2010). Hence, we report the posterior estimates of the connectivity parameters that optimally explain the group mean and difference effects across HC and SZ subjects.

#### 2.5.3 Visualization of network dysconnection

Following PEB analysis, we constructed a binary *n* × *n* matrix for each network (containing *n* subnetworks), where 1’s denoted the connections contributing to group difference effects. Then, the nodal degree for each subnetwork was computed as the sum of the corresponding row and column entries in this binary matrix. These degrees were normalized (i.e. divided) by the maximum degree in an n-node directed network (i.e. 2n-1). For visualization purposes, the normalized degrees were set as the radii of spheres centered on the peak coordinates of the subnetworks in MNI space. The results were illustrated using the BrainNet Viewer toolbox (Xia et al., 2013). Hence, a larger sphere marks further dysconnection of a subnetwork within its pertinent network.

### 2.6 Cognitive relevance of effective connectivity in SZ

After DCM analysis, we asked whether the EC profile of these seven large-scale networks could explain the cognitive performance of the SZ patients. Therefore, we used canonical correlation analysis (CCA) (Hotelling, 1936) to identify linear relationships between the strengths of EC parameters (*X*) and the MCCB scores of the patients (*Y*), after adjusting for age, gender and medication dosage (as CPZ equivalents). To decrease overfitting, we used an ensemble feature selection method (based on forward selection and backward elimination) to restrict the number of variables compared to the sample size. In the following, we outline the details of this procedure and the fundaments of the CCA model.

#### 2.6.1 Canonical correlation analysis

CCA is a multivariate statistical method that identifies the sources of common variation in *two* (usually high-dimensional) sets of variables, such that the identified patterns offer a compact description of *many-to-many* relations. Hence, CCA “opens interpretational opportunities that go beyond techniques that map *one-to-one* relations (e.g., Pearson’s correlation) or *many-to-one* relationships (e.g., ordinary multiple regression)” (Wang et al., 2020). Especially, in the era of “big data” neuroscience, CCA is being efficiently used to chart the links between brain, behavior, cognition, genes and disease (Calhoun and Sui, 2016; Correa et al., 2010; Mihalik et al., 2022; Mohammadi-Nejad et al., 2017; Smith et al., 2015; Wang et al., 2020; Xia et al., 2018; Zhuang et al., 2020).

Mathematically, CCA is designed to maximize the correlation between linear combinations of two datasets *X* and 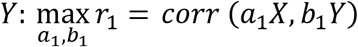. Here, *X_s×q_* contains the *q* connectivity parameters for s subjects, and *Y_s×p_* holds their MCCB scores in *p* domains^11^. As such, *r*_1_ is called the first *canonical correlation*; *a*_1_ and *b*_1_ are the first *canonical weights*; and (*u*_1_ = *a*_1_*X,v*_1_ = *b*_1_*Y*) constitute the first pair of *canonical variates.* There could be additional canonical variates (*u_i_,v_i_*) corresponding to *r*_2_,*r*_3_,…,*r*_min_(_*q,p*_) To test for the number of significant canonical correlations, Wilks’ Λ-statistic is usually used (Rencher, 2002):

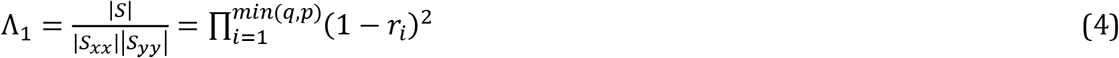

where 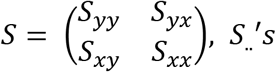 are sample covariance matrices, and Λ_1_ is distributed as Λ_*q,p,s*–*p*_. For Λ_1_ ≤ Λ_*α*_ ^12^, one would reject the null hypothesis of no linear relationship between the *x*’*s* and the *y*’*s* (i.e., *H*_0_:Σ_*yx*_ = 0). This is more evident from the second equality in Eq. 4, which shows that if one or more of the 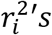 is large, then Λ_1_ will be small^13^. In practice, F- and χ^2^-approximations of Wilks’ Λ are commonly used for statistical inference (Bartlett, 1947; Rao, 1952).

Based on previous rigorous studies (Helmer et al., 2020; Leach and Henson, 2014; Yang et al., 2021), constructing reliable CCA models requires that the number of samples be at least nine to ten times the number of features. In the current dataset, cognitive scores were available for a total of 117 subjects, whereas there were 470 EC features. Hence, a suitable feature selection procedure was required to restrict the number of variables entering the predictive model. Subset selection in CCA can be performed by the same methods used in multivariate regression (Rencher, 2002). Hence, we used the *forward selection* and *backward elimination* procedures. Briefly, first a useful subset of the EC parameters (*x*’*s*) was identified using forward selection, to linearly predict the MCCB scores (*y*’*s*). Then backward elimination was applied to the *y*’*s*, pruning those that did not contribute significantly to predicting the selected *x*’*s*, in a linear model. The mathematical details are briefly revised next.

#### 2.6.2 Feature selection

We used forward selection on the EC parameters followed by backward elimination on the MCCB scores (Rencher, 2002), which we collectively call feature selection^14^ here. The forward selection method works as follows: At the first step, for each *x_j_* and (all) the *y*’*s*, a dedicated CCA model is constructed. After calculating Λ(*x_j_*) for each *j* (from Eq. 4), the variable with minimum Λ(*x_j_*) is selected and denoted as *x*_1_; this is the variable that best predicts the *y*’*s* just by itself. At the second step, we seek the variable yielding the smallest *partial Λ* – adjusted for the first chosen variable – given by:

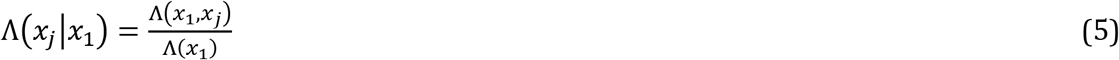

which is calculated for each *x_j_* ≠ *x*_1_. As such, the variable that minimizes Λ(*x_j_*|*x*_1_) is selected and denoted as *x*_2_. Similarly, after *m* variables have been selected, partial Λ assumes the following form for the next step:

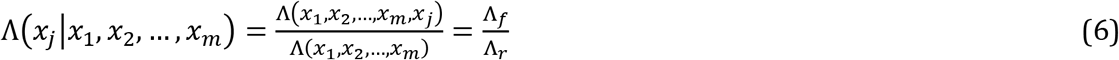

At this step, one would choose the *x_j_* that minimizes partial Λ in Eq. 6, which is distributed as Λ_*p*,1,*s*–*m*–1_ (convertible to *F_p,s–m–p_*). Notably, the second equality in Eq. 6 reinforces the fact that partial Λ is the ratio of Λ-statistics between the full(er) CCA model (containing *x_j_*) and the reduced CCA model (missing *x_j_*), denoted by Λ_*f*_ and Λ_*r*_, respectively. This selection procedure continues until the step at which the minimum partial *Λ* exceeds a predetermined threshold, or equivalently the associated partial *F* falls below a preselected value^15^. In practice, to the keep the sample to feature ratio above nine (Helmer et al., 2020; Leach and Henson, 2014; Yang et al., 2021), the feature selection was stopped earlier.

As for the backward elimination on the MCCB scores (*y*’*s*), we started with a full model (containing all the *y*’*s*, and the *x*-subset from forward selection) and deleted redundant *y*’*s* based on partial Λ. That is, at the first step we removed the *y_j_* that maximized

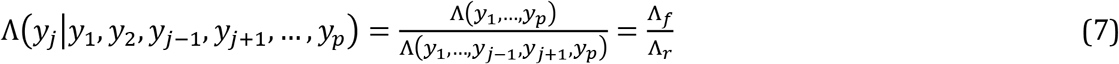

which is distributed as *Λ_1,*q*′*,s*–*p*–*q*′_* (convertible to *F_q′,s–p–q_*), with *q′* denoting the cardinality of the *x*-subset. This single variable removal procedure continued with the remining *y* variables, until a step at which the largest partial Λ became significant (at *α* = 0.05), indicating that the corresponding *y* was not redundant in the presence of its fellows. So, through feature selection, we ended up with a small subset of relevant *x*’*s* and *y*’*s* for the CCA model.

In practice, to render the model resistant to small changes in the data, we implemented an *ensemble* version of the above feature selection procedure (Bin et al., 2016; Saeys et al., 2008). That is, we used **b**ootstrap **agg**regat**ing** (aka bagging) (Breiman, 1996; Saeys et al., 2008). Specifically, 200 bootstrap resamples were generated and the feature selection procedure was applied to each resample, selecting a maximum of 5% of the variables at a time. The outputs of different feature selectors were then aggregated by computing the *frequency of inclusion* for each feature, as the proportion of times it had been selected by different feature selectors, which can range from 0 (never selected) to 1 (always selected) (Bin et al., 2016). The top few EC parameters and MCCB scores entered the final CCA model to compute the canonical correlations, weights and variates.

#### 2.6.3 Generalizability of the model

We assessed the generalizability of the linear (CCA) model using k-fold cross-validation (k=5). That is, the samples (i.e. subjects) were randomly partitioned into k subsets. The CCA model was constructed based on k-1 training folds, using the feature selection procedure outlined above. The resulting canonical weights were then used to compute canonical variates and correlations for the remaining (test) fold. In a circular fashion, each of the k folds served as the test fold exactly once. The whole cross-validation procedure was repeated 100 times, each time with a new random partitioning. As such, the out-of-sample canonical correlations were computed to test the generalizability of the linear brain-behavior model.

## 3 Results

### 3.1 Impaired cognition in SZ

Statistical tests revealed that SZ patients have impaired cognitive performance, compared to HCs, in all seven domains of the MCCB tests and in composite scores (adjusted p-values were <0.001, 0.027, 0.043, <0.001, 0.034, <0.001, 0.0362 and <0.001, respectively). Fig. 2-A summarizes these results. The box-plots show 95% confidence intervals around the mean of the test scores in red, and one standard deviation (STD) of the scores in blue/gray for HC/SZ subjects. The scores are scattered over the boxes. Specifically, the mean ± STD of the composite score for SZ patients was 33.4 ± 12.6, whereas HC subjects achieved 50.1 ± 8.2. The correlation of the subjects’ scores in different domains is illustrated in Fig. 2-B. It is apparent that, speed of processing, attention/vigilance, working memory, and reasoning/problem solving were more correlated with each other, while social cognition was the least correlated with the other (nonsocial) domains.

### 3.2 Limited FC changes in SZ

Fig. 3-A illustrates the FC patterns averaged over subjects of each group. These average patterns show the expected modular organization within sensory systems and default mode components, as well as anticorrelation between them (Chang and Glover, 2010; Fox et al., 2005; Shirer et al., 2012). Fig. 3-B depicts group differences in FC (SZ-HC), adjusted for age and gender. The colored entries reflect – *sign*(*t* – *statistic*) × log (*q_FDR_*) for the significant differences (*q_FDR_* < 0.05). From the 1225 unique FC entries, 11% were significantly different between the two groups after accounting for age and gender. As such, HCs had stronger correlation in the sensory regions and pronounced subcortical-sensory anticorrelation (than SZ patients), as reported in (Damaraju et al., 2014). However, when the regression model was adjusted for the antipsychotic medication dosage - besides age and gender - the between-group FC effects diminished further. In this case, only 18 connections (i.e., about 1% of all the functional connections) showed diagnostic effects (Fig. 3-C). The ratio was the same when considering only within-network functional connections (3/210 ≈ 0.01).

### 3.3 Numerous EC alterations in SZ

Spectral DCM was used to estimate EC between the subnetworks of each large-scale network, per subject. Fig. 4 contains representative DCM results. In Fig. 4-A, the predicted and estimated (i.e. observed) power spectral densities are plotted for the constituent subnetworks of the SM network, for an exemplar subject. The subnetworks *per se* are visualized in Fig. 4-B. The effective connections that can best predict the subnetwork spectral densities for this subject are demonstrated in Fig. 4-C. This bar plot shows the posterior expectations of the EC parameters and their estimated uncertainties (as 95% credible intervals). Positive and negative connections denote excitatory and inhibitory effects respectively; except for self-connections (*A_ii_*), which are by design inhibitory, and encoded as log scaling parameters. That is, self-connections can be converted to units of Hz using −0.5 * exp (*A_ii_*), where −0.5 Hz is the prior expected value for the self-connections. This negativity constraint ensures self-inhibition, hence stability of the dynamical system model.

**Fig. 4:**
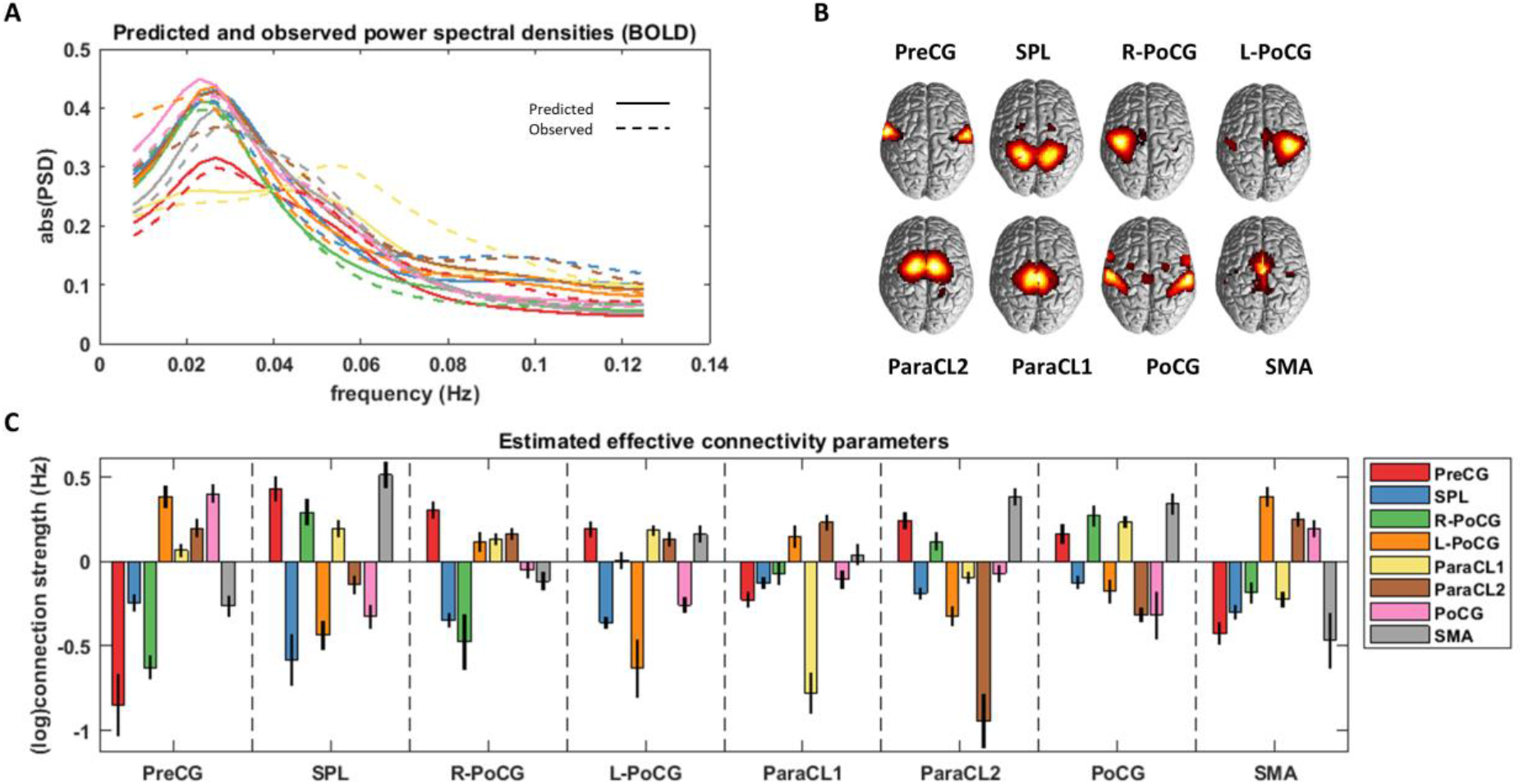
Representative spectral DCM results for a single subject. (A) The absolute values of the predicted and observed power spectral densities of the eight subnetworks in the SM network. (B) Spatial maps of the SM subnetworks. (C) The posterior expectaions (colored bars) and uncertainties (95% credible intervals; black lines) of the effective connections between the SM subnetworks. The x-labels denote the receiving ends of the directed connections, while the sources of influence are distiguished by the color codes. The extrinsic and intrinsic connections are in units of Hz and log Hz, respectively.Subnetwork abbeviations are listed in Table 2.

To assess the model fitting, the coefficient of determination (*R*^2^) was computed, which reflects the proportion of variance in the observations (herein cross spectra) that is explained by the model. The closer the value of *R*^2^ to 1, the better the model fits the data. These values were computed for each network and subject, and averaged over subjects to quantify the goodness of fit for different networks. The results are summarized in Table 3, where *R*^2^ is expressed in percentage. On average, the models explained 90±3 % of the data, across networks and subjects.

**Table 3:**
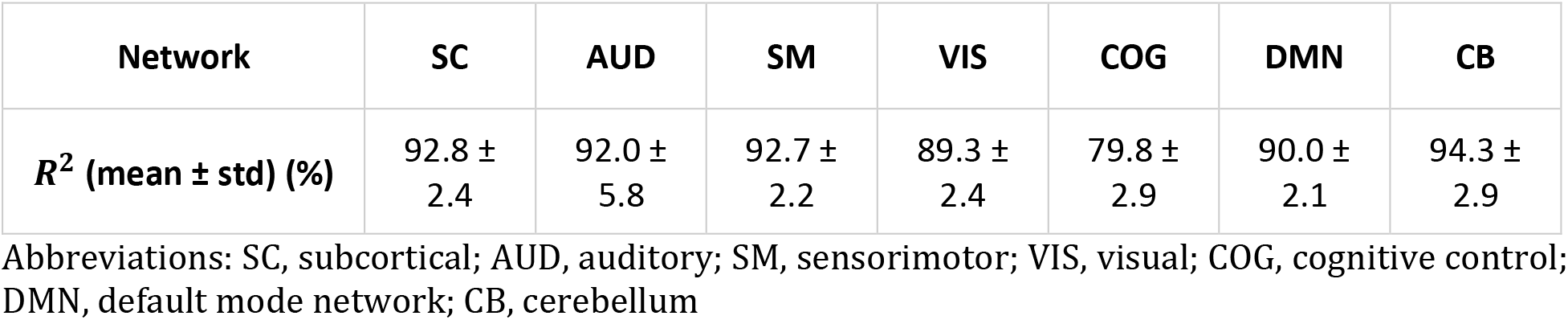
Model fitting for each network (mean ± std of *R*^2^ across subjects).

| Network | SC | AUD | SM | VIS | COG | DMN | CB |
| --- | --- | --- | --- | --- | --- | --- | --- |
| $R^2$ (mean ± std) (%) | 92.8 ± 2.4 | 92.0 ± 5.8 | 92.7 ± 2.2 | 89.3 ± 2.4 | 79.8 ± 2.9 | 90.0 ± 2.1 | 94.3 ± 2.9 |

Abbreviations: SC, subcortical; AUD, auditory; SM, sensorimotor; VIS, visual; COG, cognitive control; DMN, default mode network; CB, cerebellum

Following the first-level analysis, the posterior densities of subject-specific EC parameters were assimilated in a Bayesian GLM (i.e. PEB model) to estimate the group average of each connection strength, and the effect of diagnosis on each connection (i.e. group differences), having accounted for the effect of age, gender and medication dosage. Group differences are illustrated in Fig. 5 for each network. The plots in Fig. 5 follow the same visualization convention as the single-subject result in Fig. 4, except that here the connection strengths pertain to group (difference) effects. Furthermore, both group mean and differences are succinctly visualized as matrices in Fig. 6. Each entry *A_ij_* of these matrices denotes the expected value of group effect from subnetwork *j* to *i.* Since self-connections are log scaling parameters (as explained earlier), more positive diagonal entries denote higher regional self-inhibition in the patient group. Group-specific averages have been illustrated in supplementary Fig. S1.

**Fig. 5:**
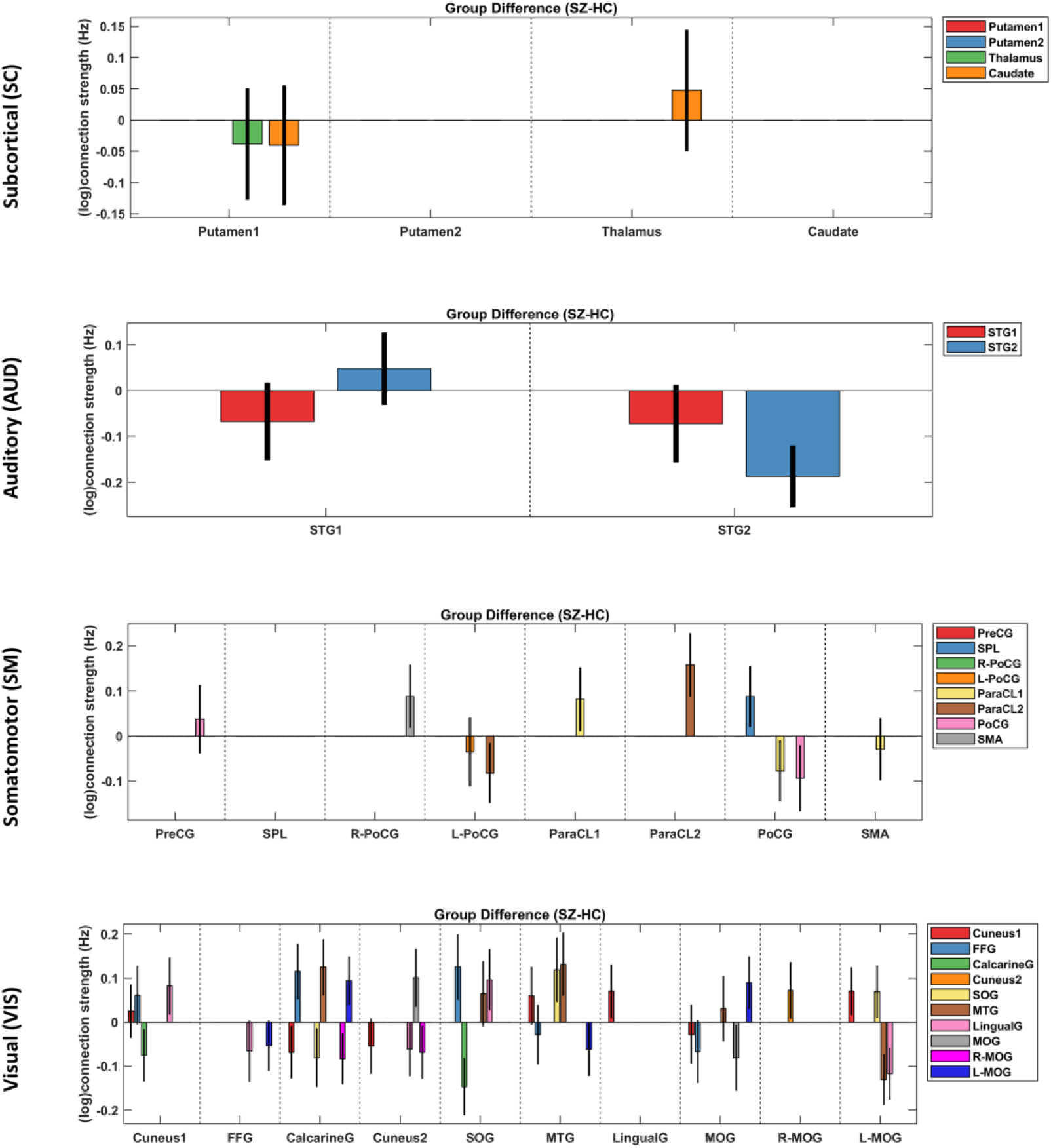
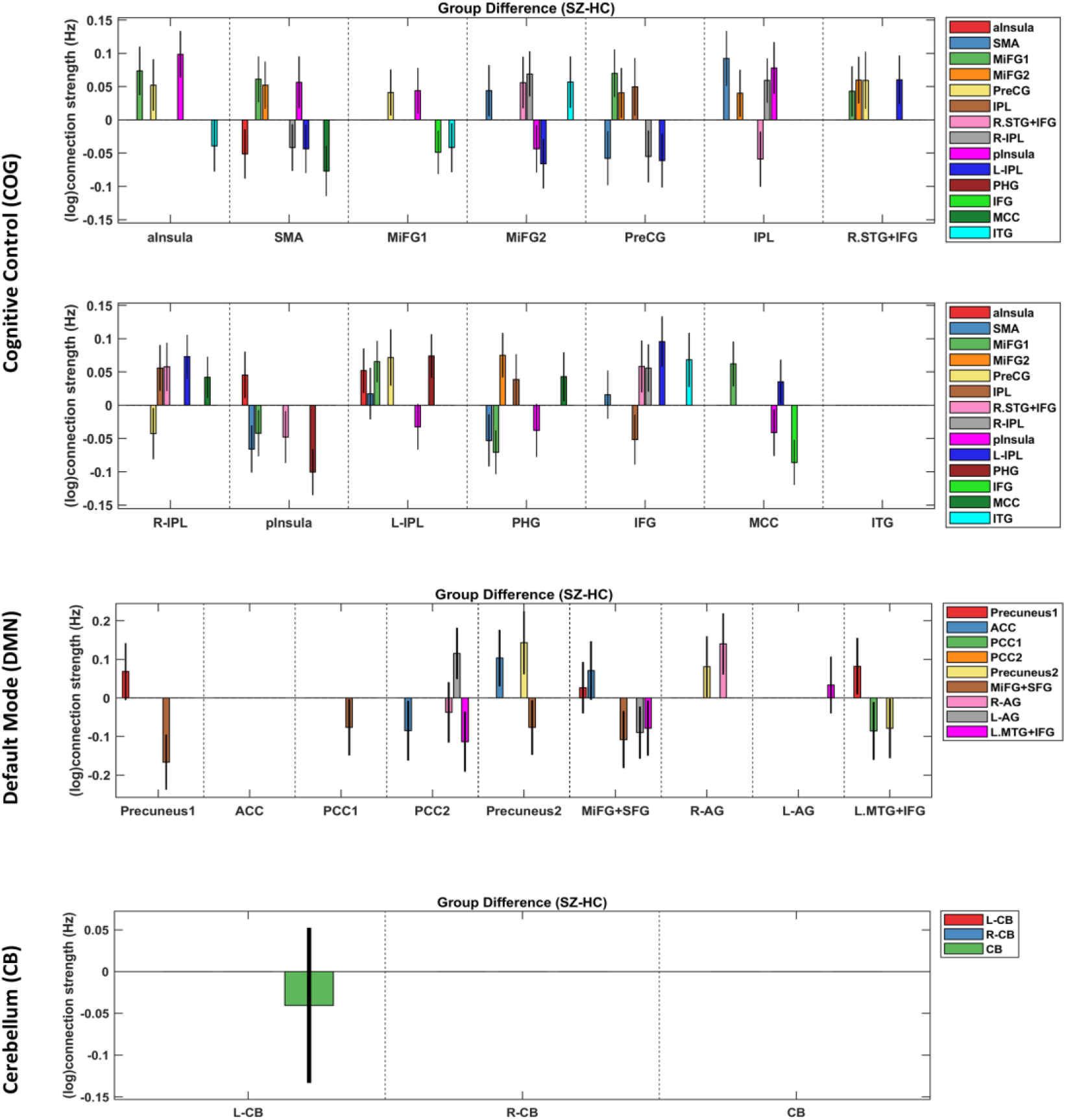
Group differences (SZ – HC) in effective connectivity. Each panel contains the posterior expectations (colored bars) and 95% credible intervals (black lines) of the effective connections between the subnetworks of one large-scale network. The x-labels denote the receiving ends of the directed connections, while the sources of inflence are distiguished by the color codes. The extrinsic and intrinsic connections are in units of Hz and log Hz, respectively. Subnetwork abbeviations are listed in Table 2.

**Fig. 6:**
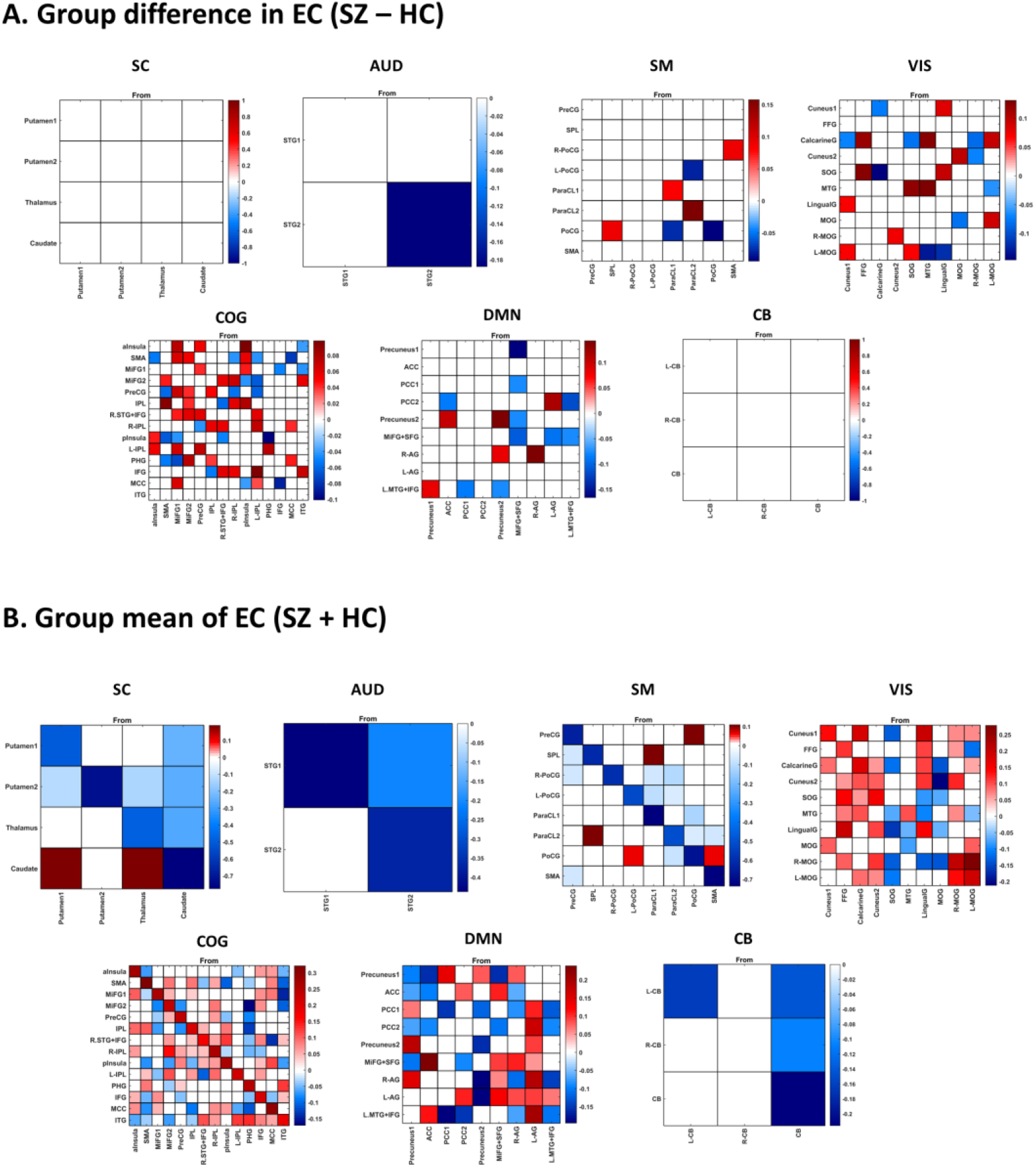
Group effective connectivity results presented in matrix format. (A) Group differences in effective connectivity (SZ – HC) between the subnetworks of each large-scale network. Each entry *A_ij_* denotes the expected value of group effect from subnetwork *j* to *i.* Only significant entries (95% credible interval not containing zero) have been colored. Diagonal entries encode inhibitory self-connections as log scaling parameters (that can be converted to units of Hz using −0.5 * exp (*A_ii_*), as elaborated in the main text). Hence, more positive diagonal entries denote higher self-inhibition effect in the patient group. (B) Group mean effective connectivity across all subjects. The format is the same as panel (A). Subnetwork abbeviations are available in Table 2.

To summarize the EC group difference results: the COG network shows the highest ratio of altered connections (33%), followed by the VIS (24%), DMN (20%) and SM (11%) networks. In the AUD network, the self-connection of (bilateral) STG2 was less inhibitory in SZ. No EC alteration was detected within the SC and CB networks, having accounted for the effect of age, gender and medication dosage. Within the COG network, 64% of the altered connections were more positive/less negative in SZ. A similar trend was seen in the VIS network (59% more positive connectivity changes in SZ). However, the opposite ratio held in the SM and DMN (57% and 69% of their altered connections were more negative in SZ, respectively). This diverse pattern of changes within multiple large-scale networks speaks to the complexity of the connectomic disorganization in schizophrenia. Remarkably, 24% of the directed influences showed SZ-related changes when analyzed with EC, whereas this ratio was 1% for the FC analysis, when the medication effect was accounted for. We shall elaborate on the implications of these findings in the Discussion.

### 3.4 Cognitive control network is the most dysconnected in SZ

Network dysconnection in SZ was further elucidated using graph theoretical methods. The normalized degrees of the subnetworks, computed from the binary matrices of significantly altered connections for each network, have been visualized on the glass brains in Fig. 7-A. The color-coded spheres are centered on the peak coordinates of the subnetworks. The size of each sphere is proportional to the extent of abnormal coupling of the corresponding subnetwork in SZ, within its associated network. As apparent by visual inspection (and the bar plot in Fig. 7-B), the COG, VIS, DMN, SM and AUD networks show considerable dysconnection in SZ.

**Fig. 7:**
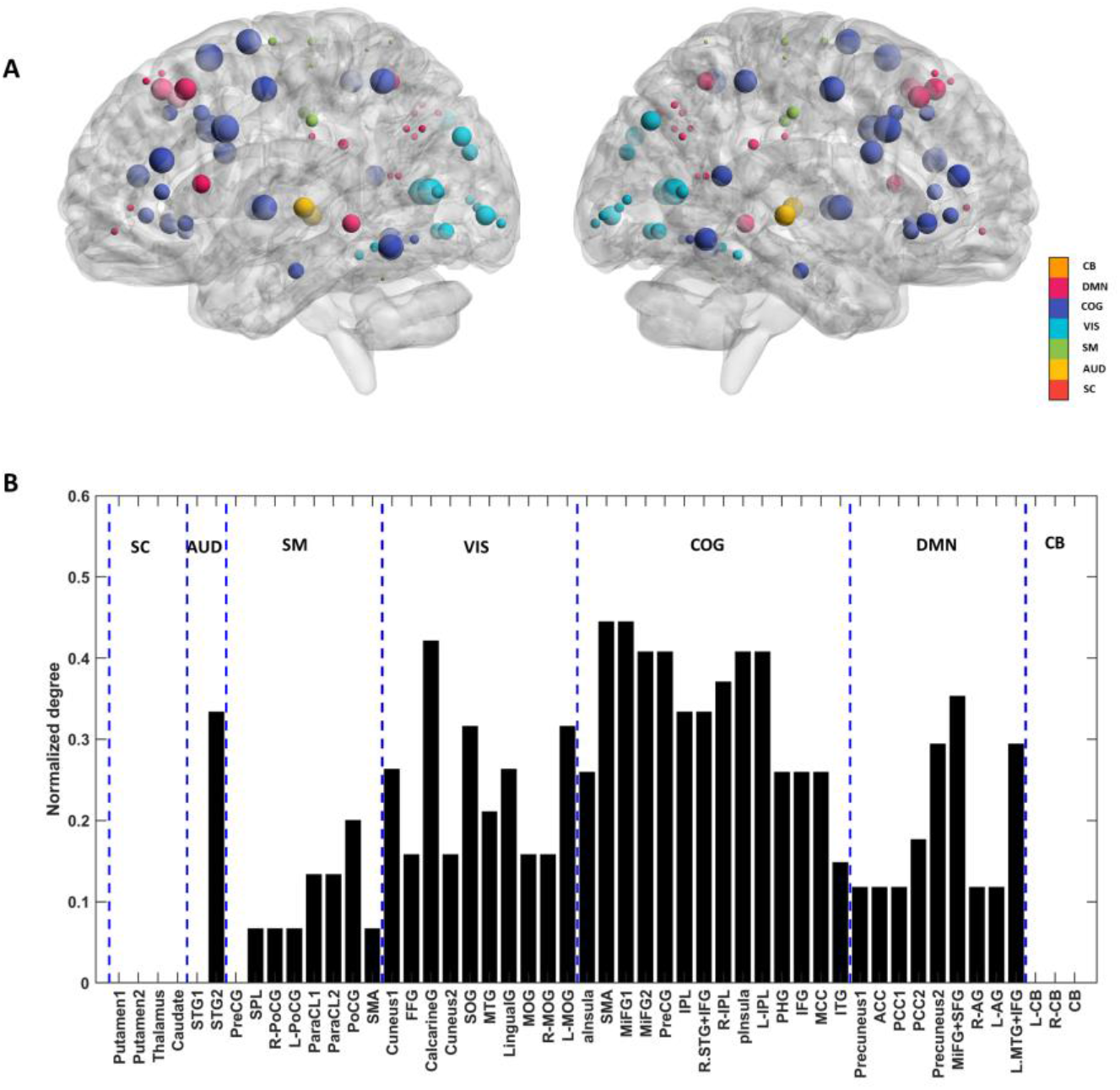
Network dysconnection. (A) The spheres are centered on the peak coordinates of the subnetworks in MNI space. The colors encode different networks. Each sphere radius is proportional to the normalized degree of the corresponding subnetwork, where the normalized degree is computed from the binary matrix of significantly altered connections (SZ – HC) from group EC analysis. The values of the normalized degrees are reported in panel (B). In short, the size of each sphere reflects the extent of abnormal coupling of that subnetwork in SZ, within its associated network.

### 3.5 Cognitive performance is related to the EC profile of SZ patients

To examine the association between the EC profile of the patients and their cognitive performance, we used canonical correlation analysis. To construct a reliable CCA model, the number of features was limited using ensemble feature selection, based on bootstrap aggregation (section 2.6.2). The resultant top features and their inclusion frequencies are listed in Table S1. The top three EC parameters included: the self-connections of SMA and ParaCL1 (in SM network) and the connection from PHG to ITG (in COG network). The inclusion frequencies of these parameters were 0.76, 0.185 and 0.18, respectively, across bootstrap resamples. Similarly, the top three MCCB cognitive traits turned out to be: social cognition, reasoning/problem solving, and working memory; with inclusion frequencies of 1.0, 0.97 and 0.93, respectively.

CCA between the selected (EC and MCCB) variables revealed one significant canonical correlation (r1 = 0.79, p < 0.001, Chi-squared = 60.05, df = 9). The corresponding (standardized) canonical weights have been plotted in Fig. 8-A. The self-inhibition of SMA was the most contributive EC parameter, followed by the self-inhibition of ParaCL1, and the excitatory connection from PHG to ITG. These three connections are positively related, based on the signs of their canonical weights. On the behavioral side, reasoning/problem solving had the highest contribution, followed by working memory and social cognition. Working memory was negatively related to the other two domains. Notably, this linear relationship was achieved with a sample to feature ratio of 58/(3+3) ≈ 9.7, in line with recent recommendations for multivariate model stability (Helmer et al., 2020; Yang et al., 2021).

**Fig. 8:**
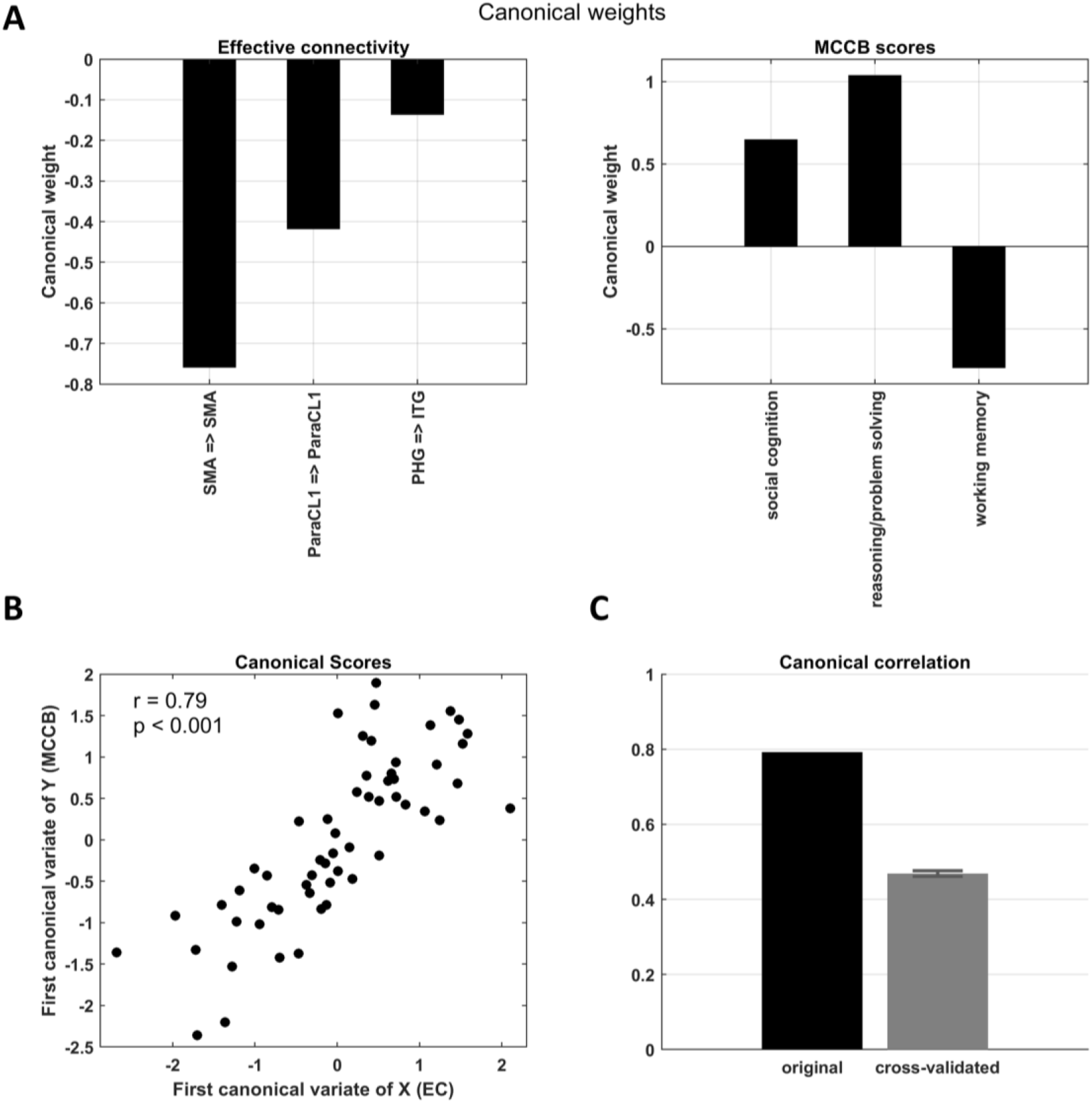
Cognitive correlates of effective connectivity, identified using canonical correlation analysis. (A) Standardized canonical weights of the selected EC features (left) and MCCB domains (right). (B) The first pair of canonical variates/scores plotted against each other. The first canonical correlation is significant (r = 0.79, p < 0.001). (C) Comparison of the original and cross-validated canonical correlations, with the latter reflecting potential generalizability of the model. Abbreviations: SMA, supplementary motor area; ParaCL1, Paracentral lobule 1; PHG, Parahippocampal gyrus; ITG, Inferior temporal gyrus.

The corresponding first pair of canonical variates/scores have been plotted against each other in Fig. 8-B. Furthermore, Fig. 8-C shows the cross-validated canonical correlation (*r_cv_*= 0.47; 0.95% confidence interval = [0.48, 0.46]), which is significant but lower than the in-sample association strength. This attenuation was anticipated based on previous studies (Helmer et al., 2020). Briefly, cross-validated estimates of canonical correlations underestimate the true values almost as much as in-sample estimates tend to overestimate them (Helmer et al., 2020). Nevertheless, the key point of this analysis is that such a linear model of brain-behavior association holds some level of out-of-sample generalizability. This is a consequence of effective feature selection and observing the requisite sample to feature ratio (section 2.6.2).

## 4 Discussion

This study investigated effective and functional connectivity between the subnetworks of seven large-scale networks of the brain, based on resting state fMRI scans from 66 SZ and 74 HC subjects. Group means and differences in EC were estimated using a parametric empirical Bayesian scheme. Group FC differences were estimated in a multiple regression model. Behavioral results from MCCB tests were analyzed separately (using permutation-based t-tests) and in conjunction with EC estimates (using canonical correlation analysis). In the following we discuss our main findings and their implications.

The analysis of behavioral data revealed significantly reduced cognitive performance in the patient group, in all seven domains of the MCCB tests and in the composite scores (Fig. 2-A). These results are consistent with the large body of literature on neurocognitive impairments in SZ (Green et al., 2019). Neurocognition typically includes: speed of processing, verbal learning and memory, visuospatial learning and memory, working memory, attention/vigilance, and reasoning/problem solving (Nuechterlein et al., 2005). Across these domains, typical SZ impairment has been reported to be 0.75-1.5 standard deviations away from that of the HC (Green et al., 2019; Heinrichs and Zakzanis, 1998; Mesholam-Gately et al., 2009). In addition to neurocognitive impairments, our results revealed significant deficits in the ‘social cognition’ of the patients. We will come back to the significance of impaired social and non-social cognition in SZ, when we discuss brain-behavior associations.

In our neuroimaging analysis, the model-based EC method identified considerably more differences between SZ and HC subjects than the descriptive FC approach did. Specifically, 24% of the investigated connections showed group difference (diagnostic) effects in the EC analysis, whereas this ratio was 1% for the FC approach, after adjusting for age, gender and medication dosage. This is mainly because, by specifying a biologically-grounded generative model (i.e. DCM) and inverting this model to fit the observations, the hemodynamic variations are effectively disentangled from the underlying neuronal dynamics, disclosing the abnormalities at the level of neural circuitry. This central difference between EC and FC (and its diagnostic consequences) may explain why EC parameters have been more informative features for diagnosis and prognosis of brain disorders in a number of studies (Brodersen et al., 2011; Brodersen et al., 2014; Frässle et al., 2020).

Notably, among the seven networks modeled using spectral DCM, the COG network turned out to be the most disturbed network in SZ, with 33% of its connections showing diagnostic effects. The other severely affected networks were the VIS, DMN and SM with 24%, 20% and 11% of their connections modulated in SZ, respectively. While most of the connections in the COG and VIS networks had become more positive/less negative in SZ, the opposite relation held for the DMN and SM networks (i.e., more negative changes in connectivity prevailed). Furthermore, in the AUD network, the bilateral STG2 subnetwork was significantly more excitable in SZ subjects. These results add to the mounting evidence for the dysconnection^16^ hypothesis of SZ, which emphasizes that “there is abnormal (rather than decreased) functional integration among brain regions in schizophrenia” (Stephan et al., 2009).

Alterations within (and across) the COG, VIS, DMN, SM and AUD networks at resting state have been reported in numerous SZ neuroimaging studies; for instance: (Bastos-Leite et al., 2015; Cui et al., 2015; Damaraju et al., 2014; Hu et al., 2017; Li et al., 2019; Uscătescu et al., 2021; Zarghami et al., 2020). However, we have not come across any prior DCM study that has modeled as many as seven large-scale networks in schizophrenia. To our knowledge, the number of regions (14 subnetworks) included in the dynamic causal modelling of the COG network is also unprecedented. This is important, because we showed that FC could not reveal the connectomic changes that EC did. Hence, the ease of conducting a large FC analysis comes at the cost of the information that can only be uncovered through model inversion of a biologically-grounded generative model.

In the SM network, we found that the self-connections of ParaCL1 and ParaCL2 were more inhibitory in SZ, which translates to lower excitability of these regions. Conversely, PoCG was more excitable (i.e. disinhibited) in the patients. In DCM framework, these (inhibitory) self-connections reflect the excitability or postsynaptic gain of neuronal populations. Computationally, these gains have been attributed to the *precision* of prediction errors (encoded by the activity of superficial pyramidal cells) ascending from lower to higher levels in cortical hierarchies, to update the predictions (encoded by deep pyramidal cells) passed down from higher to lower levels. It has been proposed that these predictions serve to explain our sensations by minimizing the prediction errors—in a theoretical framework known as predictive coding (Bastos et al., 2012; Clark, 2013; Friston, 2008; Rao and Ballard, 1999; Srinivasan et al., 1982). From the perspective of predictive coding, psychotic symptoms and failure of functional integration (i.e. dysconnection) in SZ can both be explained by aberrant precision (i.e. abnormal postsynaptic gain control) in this disorder (Adams et al., 2013; Friston et al., 2016a).

Physiologically, aberrant gain modulation in SZ has been attributed to NMDA^17^ receptor dysfunction and GABAergic^18^ abnormalities, which create excitation-inhibition imbalance and subsequently disturb the synchrony of large-scale networks (Gao and Penzes, 2015; Jardri and Deneve, 2013; O’Donnell, 2011). In the current study, altered self-connection in three subnetworks (ParaCL1, ParaCL2 and PoCG) of the SM network provided evidence for the failure of neuromodulatory gain control in these regions for SZ patients. In addition to the SM network, there was evidence for SZ-related excitation-inhibition imbalance in several other networks. Namely, bilateral STG2 in the AUD network; MTG and MOG in the VIS network; Precuneus2, MiFG+SFG and R-AG in the DMN, showed significantly altered gain control in the patient group. Still, neuromodulatory gain control in the SM network had the highest association with cognitive performance in SZ, as we will shortly revise.

Our brain-behavior analysis (using CCA) revealed that resting state EC within large-scale networks is significantly correlated with the cognitive profile of the patients. Cross-validation confirmed that such a linear association has some level of out-of-sample generalizability. To achieve a stable and generalizable CCA model, we had observed the sample to feature ratio of above nine (Helmer et al., 2020; Yang et al., 2021) using ensemble feature selection (Bin et al., 2016; Saeys et al., 2008). We shall now discuss the significance of the effective connections and cognitive domains that the feature selection procedure returned—and their contribution towards the connection-cognition model.

The top selected EC feature - by far - was the self-connection of supplementary motor area (SMA) in the SM network, which had an inclusion frequency of 0.76 over bootstrap resamples, and the largest absolute canonical weight in the CCA model. Structural and functional aberrations of SMA are well-known in the context of SZ and psychosis (Exner et al., 2006). Structurally, the volume of SMA/pre-SMA has been reported to be smaller in SZ patients, and related to their impaired implicit learning (Exner et al., 2006). Functionally, there have been recurrent reports about the reduced activation of SMA during motor and mental tasks (Crespo-Facorro et al., 1999; Guenther et al., 1994; Ortuno et al., 2005; Rogowska et al., 2004; Schröder et al., 1995) and a wide range of neurocognitive tasks in SZ (Picó-Pérez et al., 2022) and in early psychosis patients (Horne et al., 2022; Vanes et al., 2019). SMA dysfunction has also been associated with temporal processing deficit in SZ (as a measure of cognitive malfunction) (Alústiza et al., 2017; Davalos et al., 2011; Ortuño et al., 2011) and with reduced sense of agency (a common positive symptom of psychosis) (Farrer et al., 2003; Nachev et al., 2008; Wolpe et al., 2020; Yomogida et al., 2010). Our CCA analysis revealed that the self-inhibition of SMA at resting state is significantly correlated with the cognitive performance of SZ patients, above and beyond the other effective connections.

The second most contributive connection to the EC-MCCB relationship was the self-connection of ParaCL1, in the SM network. Paracentral lobule alterations have come up frequently in neuroimaging studies of SZ and psychosis. Structurally, (Borgwardt et al., 2007) found that patients with first episode psychosis and individuals (with at-risk mental state) who later became psychotic both had smaller gray matter volume in the ParaCL region, compared to HCs. Similarly, twins with schizophrenia were reported to have less ParaCL cortical volume than their nonpsychotic cotwins (Borgwardt et al., 2010). Evidence for behavioral association of ParaCL function in SZ includes the study of (Gao et al., 2022), who reported that fractional amplitude of low-frequency fluctuations (fALFF) in the ParaCL region is decreased in patients with SZ, and is associated with their clinical characteristics. Regarding cognitive relevance, reduced ALFF in the ParaCL was reported to be negatively correlated with the (impaired) speed of processing in SZ patients (Wang et al., 2019). In the present study, ParaCL excitability turned out to be both a diagnostic and a cognitively relevant connectomic feature for SZ, highlighting the implication of this region in the pathophysiology of SZ and in the psychopathological consequences.

The third contributing variable to the connection-cognition CCA model was the effective connection from PHG to ITG, in the COG network. Structurally, both the PHG and ITG regions have been reported to have less gray matter volume in SZ patients than in HCs (Curtis et al., 2021; Zhuo et al., 2017). According to (Curtis et al., 2021), the thinner gray matter in the PHG of first episode SZ patients correlates with their hallucinations, processing speed, working memory, and verbal learning. Functionally, (Diederen et al., 2010) showed that auditory verbal hallucinations in SZ patients are consistently preceded by deactivation of the PHG. Aberrant activity and connectivity patterns including the PHG and ITG have been frequently associated with working memory deficits in SZ (Chatterjee et al., 2019; Kim et al., 2009; Meyer-Lindenberg et al., 2001). We found that the excitatory influence of the PHG component on the ITG subnetwork at resting state is significantly correlated with the working memory performance, social cognition, and reasoning/problem solving capabilities of SZ patients.

Notably, the cognitive traits that correlated most consistently with EC included both social and nonsocial (i.e., working memory and reasoning) aspects of cognition in SZ. These three cognitive domains had nearly perfect inclusion frequencies during ensemble feature selection (Table S1). Nonsocial cognitive (aka neurocognitive) alterations and their neural substrates have been a major focus of SZ studies for many years. Conversely, social cognition has more recently been attended to, and proposed as a Research Domain Criteria (RDoC) domain (Cuthbert and Insel, 2013; Gur and Gur, 2016).

Social cognition broadly encompasses the mental operations needed to perceive, interpret and process information for adaptive social interactions (Green et al., 2019). Examples include emotion processing, social perception and mentalizing (aka theory of mind, ToM), in which SZ patients have consistently performed poorly (compared to HCs) with large effect sizes (Savla et al., 2013). We found significantly impaired social cognition in SZ patients (Fig. 2-A), in line with previous reports (Burns, 2006; Green et al., 2019; Savla et al., 2013). Moreover, in our brain-behavior analysis, social cognition was the only cognitive trait that was consistently selected as neurally-relevant across all data resamples (with inclusion frequency = 1). Social cognition was also the least correlated with the other (nonsocial) cognitive scores of the subjects (Fig. 2-B), which speaks to the originality of the information conveyed by the social aspect.

Overall, there is growing evidence that both social and nonsocial cognitive deficits are core features of SZ, which exist at the illness onset, cannot be explained by positive symptoms or antipsychotic medication effects, are relatively stable over the course of illness, and are detectable at lower levels in unaffected relatives of the patients and in prodromal or other high-risk samples (Burns, 2006; Green et al., 2019; Lee et al., 2015; McCleery et al., 2014). For instance, (Aiai et al., 2017) found significant differences in the working memory and reasoning/problem solving capabilities of SZ patients’ fathers compared to matched HCs. There is also evidence for heritability of emotion identification efficiency (a social cognitive trait) in SZ (Gur et al., 2007a; Gur et al., 2007b). In the present SZ sample, we found that social cognition, working memory and reasoning/problem solving are strongly correlated with the EC profiles of the patients.

Revisiting our brain-behavior results, we found that eight out of the top eleven cognitively-relevant EC features were situated in the COG network (Table S1), which is an interesting finding. Nevertheless, more remarkably, the top two cognitively-relevant connections were not in the COG network, but in the SM network; namely, the self-connections of SMA and ParaCL1. We already discussed the implications of aberrant postsynaptic gain control and excitation-inhibition imbalance in SZ, from the perspective of predictive coding. The fact that abnormal gain modulation in the sensorimotor network is so relevant to the cognitive performance of the patients also speaks to the close relationship between sensorimotor processing deficits and cognitive impairments in SZ (Fritze et al., 2022; Kumari et al., 2008; San-Martin et al., 2020).

The well-known sensorimotor and sensory gating deficiencies in SZ patients (and their unaffected relatives) (Braff et al., 1992; Braff and Light, 2005) have been attributed to their inability to filter out irrelevant external and internal stimuli, which may lead to misperceptions, sensory flooding, distractibility, disorganized thinking and cognitive fragmentation (Braff and Light, 2005; Dawson et al., 2000). A recent study on sensorimotor control in SZ showed that the modulation of cortical excitability and inhibition (during a visuomotor task) is impaired in SZ patients, and that their inferior ‘visuomotor’ performance is correlated with their ‘attention’ scores (Carment et al., 2019). Although SZ has often been regarded as a primarily cognitive disorder, it has been argued that its “symptoms may in fact rather reflect cumulative cascade impairments originating in sensory and perceptual dysfunctions, in combination with failed integration between lower- and higher-order processes” (Kaufmann et al., 2015). The considerable proportion of diagnostic connections that our EC analysis revealed within the VIS, SM and AUD networks (Fig. 6-A) highlights the crucial role of sensory/sensorimotor processing deficits in the pathophysiology of SZ. Moreover, the fact that the top cognitively-relevant effective connections included the cortical excitabilities of SM regions speaks to the importance of sensorimotor-cognition relationship in SZ.

We conclude by mentioning the limitations of the present study and some future directions of research. First, we restricted the dynamic causal modeling to *within*-network influences, for computational reasons. Estimating (causal) interactions *between* large-scale networks can enhance our understanding of the global functional organization in the brain, which has been reported to be associated with behavior and cognition in health and disease (Espinoza et al., 2019; Lacy and Calhoun, 2019; Salman et al., 2019; Tsvetanov et al., 2016; Vidaurre et al., 2021; Zhou et al., 2018). Specifically, recent reformulations of DCM (as a Bayesian linear regression model) (Frässle et al., 2017; Frässle et al., 2021) are apt for EC analysis on large networks of several hundred regions, which would facilitate both within- and between-network inferences.

Moreover, our EC results did not show SZ-related alterations among the subnetworks of the SC or CB network (although FC suggested reduced subcortical-sensory anticorrelation in SZ, similar to (Damaraju et al., 2014)). The literature contains numerous reports on the structural and functional alterations of SC regions in SZ, and their cognitive association (Andreasen et al., 1998; Carlsson and Carlsson, 1990; Fan et al., 2019; Kambeitz et al., 2014; Koshiyama et al., 2018a; Koshiyama et al., 2018b; Patterson, 1987). The implication of CB in SZ (especially through the cerebello-thalamo-cortical circuitry) has also been extensively investigated (Andreasen et al., 1998; Andreasen and Pierson, 2008; Bernard and Mittal, 2015; Brady et al., 2019; Cao and Cannon, 2019; Yeganeh-Doost et al., 2011). In the present work, the group difference in EC between two subcomponents of the CB network was explained away by the effect of medication dosage (Fig. S2). We speculate that a different or more refined parcellation of the SC and CB networks prior to EC analysis might facilitate the identification of potentially altered EC within these networks. Subcortical-cortical causal influences in SZ have also been the focus of several small-DCM studies (Csukly et al., 2020; Sabaroedin; Sabaroedin; Sabaroedin et al., 2021), which can greatly benefit from the whole-brain modeling approach furnished by recent reformulations of DCM (Frässle et al., 2021).

We accounted for the effect of antipsychotic medication dosage (in terms of CPZ equivalents) in the group (FC and EC) analyses, and in the brain-behavior (CCA) model, alongside the other covariates of age and gender. While the effect of the medication dosage covariate *per se* was not significant (in neither connectivity analysis), including the medication dosage as a covariate explained away a good number of the diagnostic connections in both FC and EC analyses (Fig. 3 for FC; Fig. 6 and Fig. S2 for EC), which highlights the confounding effect of this factor. To systematically study the neuromodulatory role of medication effect versus the core pathophysiology of SZ, longitudinal studies on drug-naïve patients (Anticevic et al., 2015; Hadley et al., 2016; Towlson et al., 2019), especially if analyzed in the framework of dynamic causal models, would be invaluable.

Another interesting avenue could be investigating the predictive power of EC versus FC features for HC/SZ classification, and for prediction of the clinical symptoms and behavioral scores. There have been several reports of the superiority of EC features (over FC) for the classification of SZ (Brodersen et al., 2014) and aphasic (Brodersen et al., 2011) patients from HCs, and for prediction of individual clinical trajectories in depression (Frässle et al., 2020), based on small DCMs. In the present study (including seven large-scale networks), the diagnostic effects identified based on EC turned out to be far more pronounced than FC differences. Hence, we speculate that a combination of generative modeling and discriminative classifiers (aka generative embedding) would increase the accuracy and generalizability of the predictor models for SZ, beyond the more conventional FC-based predictors.

## 5 Conclusions

This study investigated resting state effective and functional connectivity between the subnetworks of seven large-scale networks of the brain, in 66 SZ and 74 HC subjects. Behaviorally, SZ patients showed inferior cognitive performance in all seven domains of the MCCB cognitive tests. EC analysis revealed that a remarkable one third of the effective connections (among the subnetworks) of the COG network have been pathologically modulated in SZ. Further dysconnection was identified within the VIS, DMN and SM networks, with 24%, 20% and 11% of their connections altered in SZ, respectively. The bilateral STG2 subnetwork in the AUD network was also more excitable in the patient group. Notably, EC uncovered considerably more diagnostic connections than FC did (24% versus 1% of the connections showed group differences in EC and FC analyses, respectively). To study the neural correlates of (impaired) cognition in SZ, we conducted a canonical correlation analysis between the EC parameters and the MCCB scores of the patients. This analysis revealed that the self-inhibitions of SMA and ParaCL1 (in the SM network) and the excitatory connection from PHG to ITG (in the COG network) are significantly correlated with the social cognition, reasoning/problem solving and working memory capabilities of the patients. Future research can investigate the potential of whole-brain EC parameters as biomarkers for (diagnosis and prognosis of) SZ and other brain disorders, and for neuroimaging-based cognitive assessment.

## Acknowledgements

P.Z. is supported by core funding awarded by Wellcome to the Wellcome Centre for Human Neuroimaging (Ref: 203147/Z/16/Z). A.R. is supported by the Australian Research Council (Refs: DE170100128 and DP200100757) and Australian National Health and Medical Research Council Investigator Grant (Ref: 1194910). A.R. is also a CIFAR Azrieli Global Scholar in the Brain, Mind & Consciousness Program.

The data analyzed in this study were obtained from the COllaborative Informatics and Neuroimaging Suite Data Exchange tool (COINS; https://coins.trendscenter.org). Data collection was performed at the Mind Research Network, and funded by a Center of Biomedical Research Excellence (COBRE) grant 5P20RR021938/P20GM103472 from the NIH to Dr. Vince Calhoun. The group ICA templates from (Allen et al., 2014) are publicly available at http://trendscenter.org/data.

## Author Contributions

**Tahereh S. Zarghami:** Conceptualization, Methodology, Analysis, Interpretation, Writing - Original Draft. **Peter Zeidman:** Methodology, Software, Interpretation, Writing - Review & Editing. **Adeel Razi:** Methodology, Software, Interpretation, Writing - Review & Editing. **Fariba Bahrami:** Methodology, Interpretation, Writing - Review & Editing, Supervision. **Gholam-Ali Hossein-Zadeh:** Methodology, Interpretation, Writing - Review & Editing, Supervision.

## Data Availability

The data analyzed in this study are available in publicly shared databases, which are listed in the Acknowledgements section.

## Footnotes

1 A “mode” is a distributed functional pattern or an intrinsic network, such as the default mode. It is contrasted against a “node” which is a local anatomical region, in the terminology of (Friston et al. (2014b)).

2 The Measurement and Treatment Research to Improve Cognition in Schizophrenia (MATRICS) Consensus Cognitive Battery (MCCB) tests (Nuechterlein et al. (2008); Kern et al. (2011); August et al. (2012)).

3 http://fcon_1000.projects.nitrc.org/indi/retro/cobre.html

4 Diagnostic and Statistical Manual of Mental Disorder, 4th edition (DSM-IV)

5 https://www.fil.ion.ucl.ac.uk/spm/software/spm12/

6 The Measurement and Treatment Research to Improve Cognition in Schizophrenia (MATRICS) initiative was designed to support the development of psychopharmacological agents to improve cognition in schizophrenia.

7 http://trendscenter.org/software/gift/

8 The artefactual components (including physiological, head motion and imaging artifact components) were identified and eliminated in (Allen et al., 2014), leaving 50 reproducible functional subnetworks. The peak coordinates of these subnetworks (herein used as spatial priors) are listed in supplementary Table 1 of (Allen et al., 2014).

9 Optimal parameters offer the most accurate and least complex explanation for the data.

10 Also known as evidence lower bound (ELBO) in machine learning.

11 The variables (i.e., the columns of *X* and *Y*) are z-scored to get standardized canonical weights (*a* and *b*).

12 Λ_*α*_ denotes the critical value of Wilks’ *Λ* corresponding to *α* = 0.05.

13 Λ_2_, Λ_3_ and so on are similarly defined to test the significance of *r*_2_ and succeeding *r_i_*’*s* after the first (Rencher (2002))

14 Alternatively, ‘feature combination’ can be used to restrict the number of variables prior to CCA, using e.g. singular value decomposition (Friston et al. (1995)). However, the resulting synthetic features would be less functionally interpretable than the original features (i.e., the effective connections), especially when discussing their contributions towards the final CCA model.

15 Alternatively, the stopping rule can be cast in terms of the p-value of the partial *Λ* or *F* exceeding a predetermined value – e.g., the conventional *α* = 0.05.

16 The Greek prefix ‘‘dys” means bad or ill, whereas the Latin prefix “dis” means apart (Stephan et al. (2009)).

17 N-methyl-D-aspartate (NMDA)

18 Gamma-aminobutyric acid (GABA)

